# Ancient biomolecular analysis of 39 mammoth individuals from Kostenki 11-Ia elucidates Upper Palaeolithic human resource use

**DOI:** 10.1101/2024.06.14.598638

**Authors:** Alba Rey-Iglesia, Alexander J.E. Pryor, Tess Wilson, Mathew A. Teeter, Ashot Margaryan, Ruslan Khaskhanov, Louise Le Meillour, Deon de Jager, Gaudry Troché, Frido Welker, Paul Szpak, Alexander E. Dudin, Eline D. Lorenzen

**Affiliations:** Globe Institute, University of Copenhagen, Denmark 1350; Department of Archaeology and History, University of Exeter, United Kingdom EX4 4QE; Department of Anthropology, Trent University, Canada K9L 0G2; Complex Research Institute of the Russian Academy of Sciences, Russia 364051; Kostenki Museum-Preserve, Voronezh, Russia 396815

**Author notes:** Corresponding authors: Alba Rey-Iglesia, Eline D. Lorenzen.

## Abstract

Circular structures made from woolly mammoth bones are found across Ukraine and west Russia, yet the origin of the bones remains uncertain. We present ten new mammoth radiocarbon dates from the largest circular structure at Kostenki 11-Ia, identifying two mammoth mandibles ∼200-1,200 years older than the other dated materials from the site, suggesting skeletal material from long-dead individuals was scavenged and used in the site construction. Biomolecular sexing of 30 individuals showed a predominance of females, suggesting the Kostenki mammoths are primarily from herds. We identify six mitochondrial lineages across 16 samples, showing they are not all from the same matriline. Integrating biomolecular sexing with stable *δ*^13^C and *δ*^15^N isotope analysis, we find no isotopically-differentiated resource use by females and males, providing the first analysis of foraging differences between sexes in any Late Pleistocene megafauna. Our study highlights the significance of integrating ancient biomolecular approaches in archaeological inference.

**Teaser:** Integrating ^14^C dating, ancient DNA, palaeoproteomics, and stable isotopes improves our understanding of Kostenki 11-Ia

## Introduction

Kostenki 11 is an archeological site embedded in a complex of 26 Upper Palaeolithic sites situated around the villages of Kostеnki and Borshchevo in south-western Russia (Voronezh Oblast; Fig. 1D) (1). The site is formed by several archeological layers. Five of these layers (layers Ia-V) have been well characterised (2); based on radiocarbon ages, they date from ∼40,000 (layer V) to ∼24,000 (layer Ia) calibrated years before present (cal yr BP) (1).

**Fig. 1.**
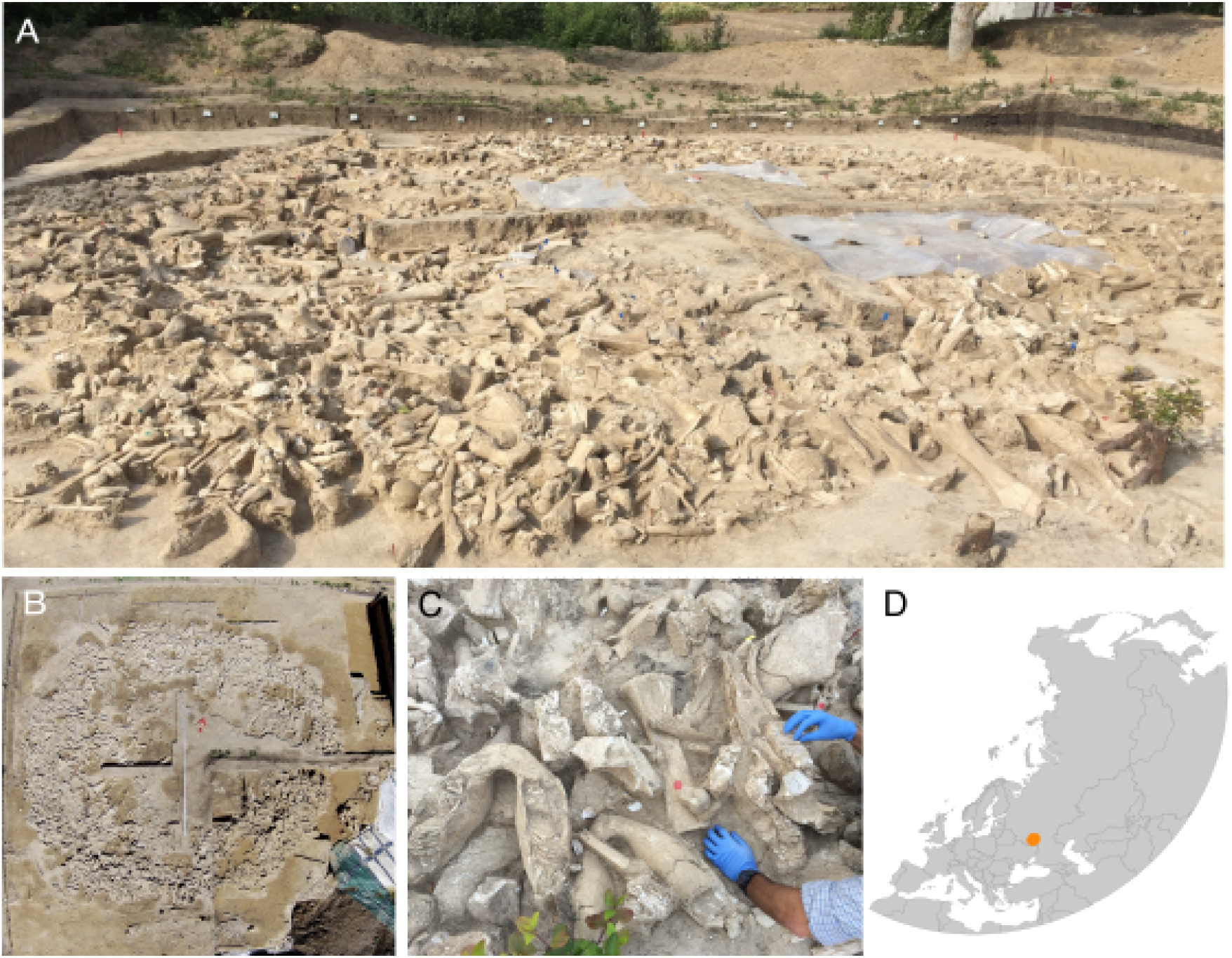
The third mammoth bone structure at Kostenki 11-Ia. (A) Panoramic image of the site taken during our sampling trip in August 2015 (Picture: E.D. Lorenzen). (B) Aerial image taken during 2015 (Picture: A.Yu. Pustovalov and A.M. Rodionov). (C) Close-up of the mammoth bones forming the structure, showing six mandibles of various sizes. (D) Map of Eurasia indicating the location of the Kostenki 11 site.

The Kostenki 11 site contains three circular mammoth bone structures in layer Ia (referred to here as Kostenki 11-Ia) (3). Circular structures made from the bones of woolly mammoths (*Mammuthus primigenius*) are known from across the North European Plain and the East European Plain. Most are found along the Desna/Dnepr River systems in present-day Ukraine and Russia (4–5), and radiocarbon dating has indicated their usage ∼22-12 ^14^C thousand years ago (kya), corresponding to ∼26-14 cal yr BP, with the majority of dates between 15.5-14 ^14^C kya, corresponding to 18.8-17 cal yr BP (4–5). The structures are usually associated with pit features potentially used for storing food or fresh bones, or discarding refuse, and indicate the past existence of open-air human settlements adapted to the steppe environment (6), while some studies also suggest these structures may have been used as ceremonial sites (7–8).

Kostenki 11 was discovered in 1951. The first mammoth-bone structure at the site was excavated during the 1960s and is now encapsulated and on display in the main building of the Kostenki Museum-Preserve (9). A second structure was discovered in close vicinity in 1970 and was only partially excavated (9). These two mammoth structures each encompass a large central (9-10 m diameter) circular structure made of mammoth bones, surrounded by storage pits.

The third mammoth-bone structure is the focus of the present study and was discovered adjacent to the Kostenki Museum-Preserve building during a survey in 2013-2014 (10). The structure is placed 20 m from the first bone structure (10). It is a large circular structure with at least three peripheral pits, and is a more sophisticated entity than the other two structures at Kostenki 11-Ia. It is larger both in size (with a central cluster of 12×11 m) and in the sheer number (n = 2,982) of mammoth bones found, with a minimum of 64 individuals identified based on the number of mammoth crania (11–12) (Fig. 1, Tables S1, S2). Previous radiocarbon dating of charcoal and faunal remains (including burnt mammoth bone) from the third structure indicate that it is one of the oldest such structures associated with modern humans yet discovered on the eastern European Plain, with dates ranging from 20,838 ± 519 to 19,514 ± 257 ^14^C yr BP, corresponding to 25,063 to 23,481 cal yr BP (3).

The mammoth bones of the structure have been investigated and described based on morphology, and extensive work has been conducted on the identification of various skeletal elements, their positioning and preservation, including the presence of animal gnawing and chewing marks and anthropogenic cut marks and notches (e.g., 10). However, additional methods are required to answer several outstanding questions, including a more reliable time frame of when and for how long Palaeolithic humans were associated with the site, and how Palaeolithic humans procured the mammoth resources used to form this structure.

Ancient biomolecules provide the toolbox necessary to further investigate the site (13). Radiocarbon dating can be used to infer when the site was in use, ancient DNA can be used for genetic sex determination of the mammoth individuals in the structure and to elucidate the phylogeographic context of the mammoths. Where ancient DNA is insufficiently preserved, ancient protein analysis may provide an alternative approach to determining genetic sex through the analysis of dental enamel. Stable isotopes contribute information on the dietary niche of mammoth individuals, and may elucidate differences in foraging ecology between females and males when used in conjunction with genetic sexing.

Due to its rich fossil record and emblematic status as a flagship species of the extinct Late Pleistocene megafauna, woolly mammoths remain one of the most well-studied species of the Eurasian mammoth steppe (e.g., 14-17). During the last glacial period ∼115-12 kya, mammoths were widely distributed across Eurasia and extended into the northern half of North America (e.g. 18). Radiocarbon dated mammoth remains have been widely investigated using ancient biomolecular approaches, including ancient DNA and stable isotopes, providing insights into their movement patterns, population histories, and palaeoecology. Indeed, the oldest nuclear genome ever sequenced is from a 1.6 million year old Siberian mammoth (19). Hence for this species, a rich panel of reference data exists with which to contextualise ancient biomolecular information from new specimens.

Woolly mammoths are believed to have had a matriarchal social structure, where adult females form groups with their offspring, and adult males leave the herd or form small, temporary bachelor groups, as this trait is shared by all extant proboscideans (20). This hypothesis is supported by morphological analysis of mammoth bone assemblages, such as the Sevsk woolly mammoth family group (Bryansk Region, Russia), which comprised a mix of females and males of different ages (21). Male-dominated sites of the Columbian mammoth (*Mammuthus columbi*), such as at the Hot Springs locality (South Dakota, USA), are believed to represent individuals outside a family group (22). However, morphological sex determination may not be reliable or indeed even possible for most fossil specimens. There have been no previous attempts to genetically sex the skeletal remains present in the mammoth structures from the central European Plain, including Kostenki. This information may provide novel insights into the origin of the skeletal remains and the hunting behaviour of prehistoric humans.

The retrieval of ancient DNA from mammoth remains has provided evidence of the biogeography of mammoths across time and space, detailing how and when mammoth genetic lineages moved across the landscape (termed phylogeography) (e.g., 14,23-25). Synthesis of the available mitochondrial genome data from mammoths shows the species can be divided into three main, genetically distinct groups (14). Although the three groups are all widely distributed across the landscape, the geographic distribution of specific subclades within each genetic group can in some cases provide information as to the geographic origin of an individual (e.g., 14,23).

Bone and dentine collagen carbon (*δ*^13^C) and nitrogen (*δ*^15^N) stable isotopes provide information on the foraging ecology of individuals. In herbivores, they provide information on the relative importance of types of plant species in the diet. The composition of plants in the diet (26) is indirectly affected by climatic and environmental factors (27–29) and can therefore be further used to make inferences about the palaeoecology of the environment inhabited by individuals/species.

In this study, we used a combined biomolecular approach of radiocarbon dating, ancient DNA, palaeoproteomics, and stable *δ*^13^C and *δ*^15^N isotope analysis, to investigate 39 woolly mammoth individuals collected from the third residential complex of Kostenki 11-Ia. Our findings represent: (i) ten new ^14^C dates, which are discussed in the context of other available direct dates from Kostenki 11-Ia, to infer the time frame of human activity at the site; (ii) combined genetic and proteomic sexing of 30 specimens, which provide indirect insights into resource use of Palaeolithic humans; (iii) mitochondrial genomes of 16 individuals, which we analysed with 147 publicly available mitochondrial genomes, to investigate the matrilineal composition of the site and the phylogeographic context of the mitochondrial sequences; and (iv) *δ*^13^C and *δ*^15^N values from 38 individuals, which we analysed with 378 available records from mammoths sampled across time and space, to detect resource partitioning between the sexes and characterise the paleoecology of the Kostenki mammoths.

## Results

### Radiocarbon dating

We radiocarbon dated nine woolly mammoth individuals from the third mammoth bone structure at Kostenki 11-Ia; one individual was dated twice, yielding a total of ten dates (Table S3). None of the woolly mammoth specimens sampled showed signs of burning, and thus burning has not altered the radiocarbon ages of these specimens.

Radiocarbon dating initially produced two dates younger than the rest of the dates new to this study, and also younger than most of the existing dates for the third mammoth structure (Fig. S1, Tables S3, S4): UCIAMS-251304 (18222; genetically sexed as female) and UCIAMS-251305 (18241; genetically sexed as female) with ages 18,080 ± 60 ^14^C yr BP and 18,870 ± 70 ^14^C yr BP, respectively. Both specimens were therefore re-dated using freshly extracted collagen taken from different parts of the molar.

Specimen 18222 failed the re-dating due to an insufficient amount of collagen. Specimen 18241 produced a second date of 20,400 ± 140 ^14^C yr BP (UCIAMS-266002), similar to the other dates new to this study, with a median age more than 1,500 years older than the first dating attempt, confirming that our two younger dates are almost certainly a result of incomplete removal of contamination during pre-treatment, exacerbated by a relatively low weight of endogenous collagen for these specimens. Difficulties with removing exogenous younger carbon contamination from collagen samples used for dating is a well-known problem with Palaeolithic material, including from other Kostenki sites (e.g., 30-32).

The eight remaining and reliable dates were combined with three previously reported dates measured on charcoal recovered from inside the third mammoth bone structure (Supplementary Text). The 11 retained dates clustered into three distinct groups (Fig. 2). The earliest group comprises two dates measured on mammoth teeth, which are notably older than all other measured dates from the site. The molars – both genetically sexed as female – were located in the outer ring of the structure (Fig. S2).

**Fig. 2.**
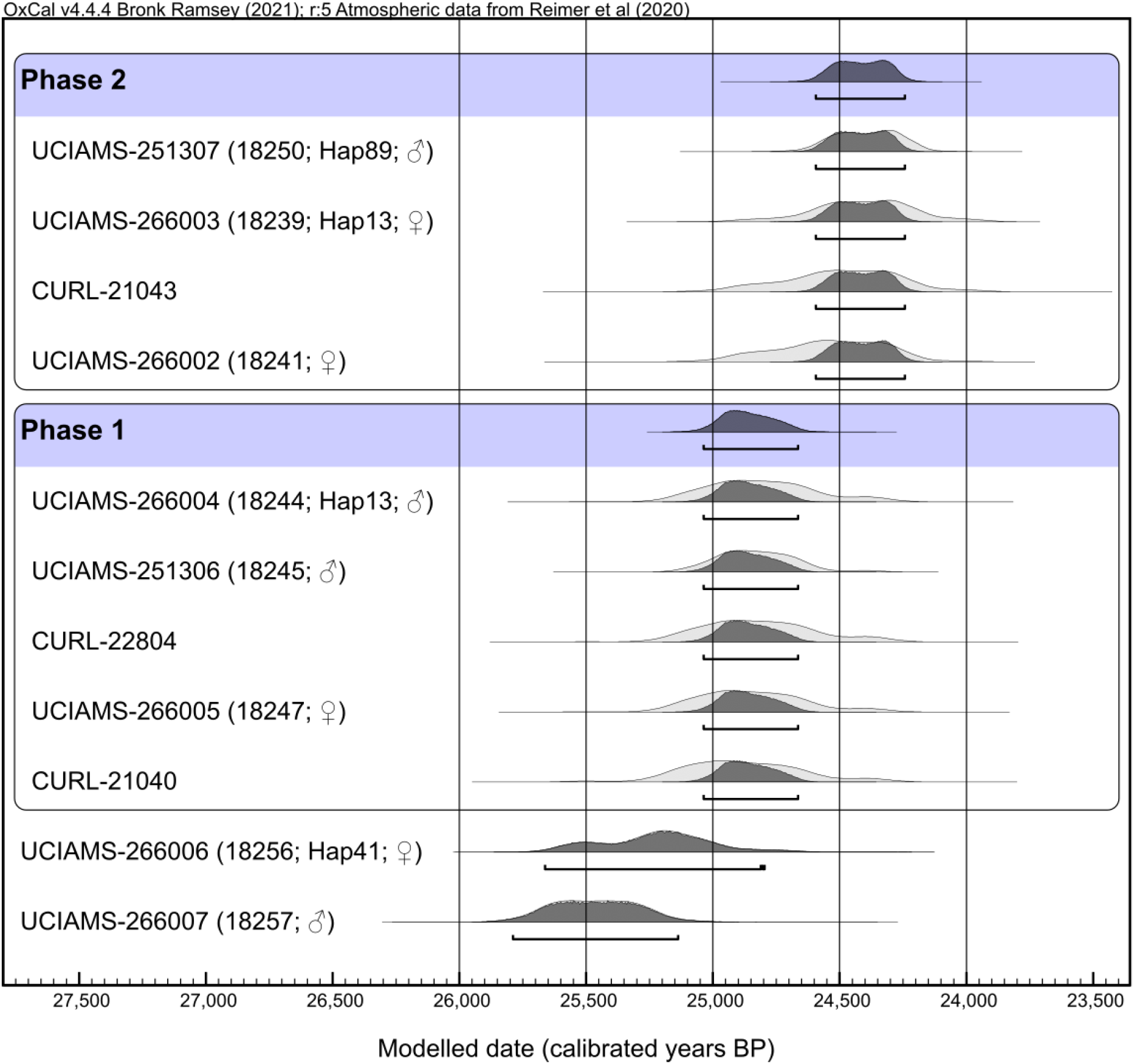
Modelled radiocarbon dates indicate two phases of human activity at Kostenki 11-Ia. (A) New (UCIAMS, all unburnt mammoth bones) and published (CURL, all charcoal) radiocarbon dates from the third mammoth bone structure. We used the ‘Combine’ function in OxCal v. 4.4 (33) to determine two discrete phases of human activity at the site: Combine Phase 1 (25,040-24,670 cal yr BP) and Combine Phase 2 (24,580-24,230 cal yr BP) at 95.4% certainty. Two bone dates were too old to be modelled with the phases (UCIAMS-266006 and UCIAMS-266007). For the UCIAMS specimens, specimen ID (abbreviated to the last five digits), mitochondrial haplotype (if available), and biomolecular sex is included. Further information on specimens and dates can be found in Supplementary Text and Tables S3, S4.

Two further groupings of dates are apparent, which we analysed using Oxcal’s ‘Combine’ function (Supplementary Text), which combines probabilities from multiple dates each giving independent information on the age of a specific sample or context (33). The method may, where appropriate, include dates measured on different materials using different methods (e.g. ^14^C, TL, OSL). Importantly, the method assumes that the activity being dated occurred within a ‘short’ time frame relative to the uncertainties associated with the dates being modelled; in this case approximately 500-1,000 years for the calibrated dates. It is therefore appropriate to use the ‘Combine’ function to model dates from the third mammoth bone structure at Kostenki 11-Ia, assuming that any phase, or phases, of activity lasted for a brief period relative to the dating uncertainty, in this case not exceeding a few hundred years at most. This assumption is well supported by the archaeological evidence at Kostenki 11-Ia, which indicates that human activity at the site actually occurred over a much briefer period or periods, lasting a few years or decades at most (3,10). We note here the difference between Oxcal’s ‘Combine’ and ‘R_Combine’ functions, the latter of which is designed to combine radiocarbon dates measured on the same artefact, and which would be inappropriate for analysing the Kostenki 11-1a dates.

When modelled with Oxcal’s ‘Combine’ function, a group of five dates measured on both mammoth teeth and charcoal (termed ‘Phase 1’, Fig. 2) have a modelled date range of 25,040 - 24,670 cal BP while the other group, comprising one date on charcoal and three on mammoth teeth (termed ‘Phase 2’, Fig. 2) have a modelled date range of 24,580 - 24,230 cal BP. There is excellent statistical agreement between the modelled Phase 1 and Phase 2 dates, respectively, with A’Combs above 140%.

The calibrated ages for our Phase 1 are remarkably similar to those indicated for Group 1 dates by Pryor et al. (3), supported by three dates new to this study. In contrast, calibrated ages for Phase 2 are several hundred years older than Pryor et al. 2020 Group 2 dates, as a direct consequence of focusing on the relatively high-precision CURL and UCIAMS radiocarbon dates, and setting aside the less-precise and likely contaminated NSKA dates (Supplementary Text).

### Genetic analysis

We estimated the endogenous DNA content of our 39 specimens by mapping the DNA sequencing reads to the nuclear genome assembly of the African savannah elephant (*Loxodonta africana*; LoxAfr4). Overall, most of the specimens had very poor DNA preservation. Endogenous content ranged from 0.002 to 26.5%, with: 23 samples having < 1% endogenous content, nine samples 1-6%, five samples with 6-9%, and two samples with an endogenous content of 18.5% and 26.5% (Table S5).

### Biomolecular sex assignments

For 23 of the 39 individuals that were DNA sequenced, which included seven of the radiocarbon dated specimens, we had enough DNA data to genetically determine sex (Table S3). We found 15 of the specimens were female and eight of the specimens were male. To increase the number of sex assignments, we applied a palaeoproteomic approach to the remaining samples.

One of the male reference specimens sexed using ancient DNA, and four of the specimens with unknown sex contained no AMELX or AMELY, making proteomic sex assignment impossible (Table S7). The proteomes recovered for these five specimens suggests that no to very little dental enamel was provided, also given the absence of other enamel-specific proteins in each of these five extracts, and that they were instead composed of dentine and/or cementum, which generally do not contain amelogenin.

In contrast, the remaining 16 extracts contained peptide matches to collagen type I (COL1A1 and COL1A2), amelogenins (AMELX and AMELY), ameloblastin (AMBN), enamelin (ENAM), collagen type 17, alpha-1 (COL17A1), matrix metalloproteinase-20 (MMP20), albumin (ALB), odontogenic ameloblast-associated protein (ODAM), and/or amelotin (AMTN), indicating the recovery of an otherwise normal Pleistocene dental enamel proteome (34–35).

We were able to correctly assign the biological sex of the remaining five specimens with known, genetically determined biological sex (3 females and 2 males; Table S7). Based on these reference specimens, we determined that a cut-off of a minimum of two unique peptides matching to AMELY and a minimum of 15 unique peptides matching to AMELX was required to confidently assign sex. We determined that five out of 11 unknown specimens represented male individuals, two represented possible male individuals (with one AMELY-specific peptide identified), two represented female individuals, and two represented possible female individuals (fewer than 15 AMELX-only peptides identified) (Table S7). All confidently identified male individuals contained at least two unique peptides overlapping amelogenin position 46 (in relation to XP_049728859.1 for AMELX and XP_049729447.1 for AMELY), where AMELX contains an isoleucine (I) while AMELY contains an methionine (M). In addition, specimens 18238, 18224, and 18259 contained unique AMELY peptides matching to other amelogenin positions (P123L; I124V; Q138H; position in reference to AMELX followed by the homologous amino acid present in AMELY, for the accession numbers listed above).

Together with the 23 mammoth specimens sexed genetically, this results in the identification of 17 (57%) females and 13 males (43%) among 30 individuals.

### Mitochondrial genomes

We generated mitogenomes for a total of 16 mammoth individuals. For three samples, we were able to compile their mitogenomes based on the shotgun sequencing data, with >1,000 DNA sequencing reads mapping to the mitogenome reference (18222, a female; 18250, a male; 18256, a female). For 14 individuals, we assembled mitogenomes with the capture approach with >900 DNA sequencing reads for each sample mapping to the mitogenome reference (Tables S3, S5).

Specimen 18256 produced a mitogenome in both the shotgun and capture approaches, which were identical at all non-ambiguous bases. We retained the captured version of the mitogenome for this sample, as it had higher coverage (715x *vs* 3.9x) and contained fewer ambiguous bases. Thus, we obtained a total of 16 mitogenomes with coverage ranging from 2.8x-715x (full sequencing statistics in Table S5).

Of the 16 new sequences, three had >20% N bases and were initially excluded from the haplotype analysis. The remaining 13 sequences were analysed with 147 publicly available mitogenomes representing all known woolly mammoth clades (Table S6). The combined data set of 160 mitogenomes represented 90 unique DNA sequences, termed haplotypes. Among the 13 Kostenki-Ia mitogenomes, we identified six haplotypes: Hap13, 41, 42, 43, 74, and 89 (Fig. 3, Tables S3, S6).

**Fig. 3.**
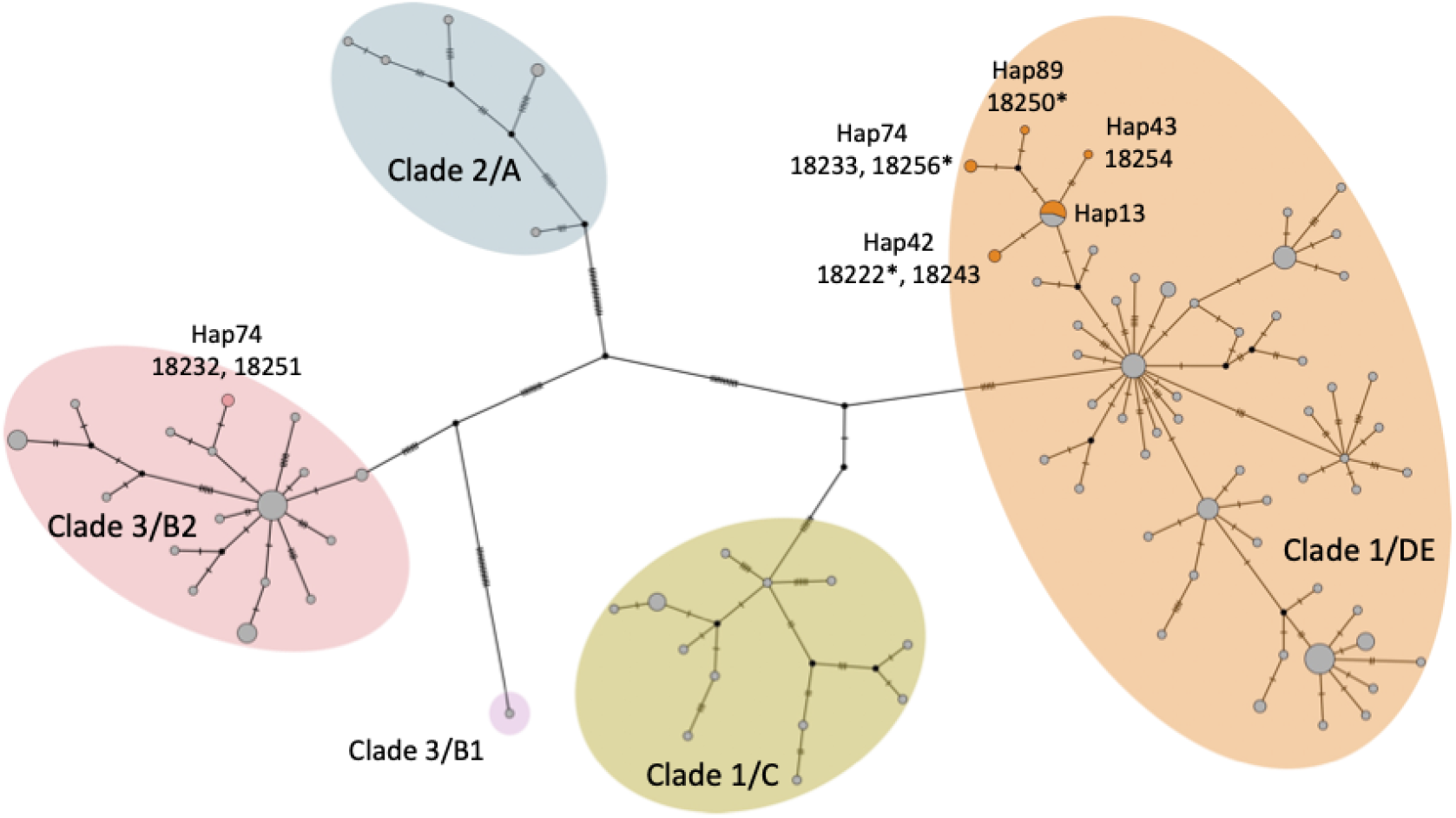
Haplotype network of 160 woolly mammoth mitogenomes. Median-joining haplotype network based on the alignment of the 13 Kostenki-Ia and 147 available woolly mammoth mitogenomes. The alignment contained 15,425 sites, of which 226 were variable. Sites with gaps or missing data were not considered. Each haplotype is indicated by a circle. The relative size of the circles represents the number of sequences of each haplotype. The new Kostenki-Ia mitogenomes are coloured in orange and pink; the haplotype numbers and Kostenki 11-Ia sample IDs discussed in the text are indicated. Previously published mitogenomes are coloured in grey. Black dots represent haplotypes not present in the data. Lines between haplotypes, which represent the evolutionary distance between two unique sequences, are not drawn to scale. The hatches on these lines show the number of substitutions between the haplotypes. Hap13 was shared among Kostenki 11-Ia specimens 18228, 18229, 18239, 18240, and 18244 (Table S3); specimens 18225, 18236, and 18255 shared haplotype Hap13, but are not included in the network analysis, as they had >20% Ns.

The three samples with >20% N bases were analysed separately to determine their haplotype affiliation (18225, 18236, and 18255). As these mitogenomes had a large amount of missing data (N’s), their inclusion in the initial analysis would have greatly reduced the number of informative sites in the alignment, thereby reducing the power of the haplotype analysis. The separate analysis of these mitogenomes indicated that all three had the most common haplotype (Hap13) of Kostenki-Ia mammoths in Clade 1/DE (Table S3).

Hap13 was present in eight of our individuals – and was also shared with the one available mitogenome from Kostenki dated to 35,170 cal yr BP (SP2401/KX176789.1; unknown Kostenki site number; 14). Hap13 was also shared with three non-Kostenki mammoths – from the Kraków Spadzista site in Poland, which are dated to ∼27,000 cal years BP (36).

The other five haplotypes present in our samples were all unique to Kostenki-Ia (Fig. 4). Haplotypes 41, 42, and 74 were each shared by two Kostenki-Ia mammoths, and haplotypes 43 and 89 were each observed in only one individual. The control region sequence was initially omitted from the mitogenome analysis due to misalignments and missing data. However, the region has a relatively fast mutation rate and is therefore more variable, and thus we analysed the d-loop specifically for the individuals sharing a haplotype, to investigate whether there were any single nucleotide polymorphisms (SNP) at this locus.

**Fig. 4.**
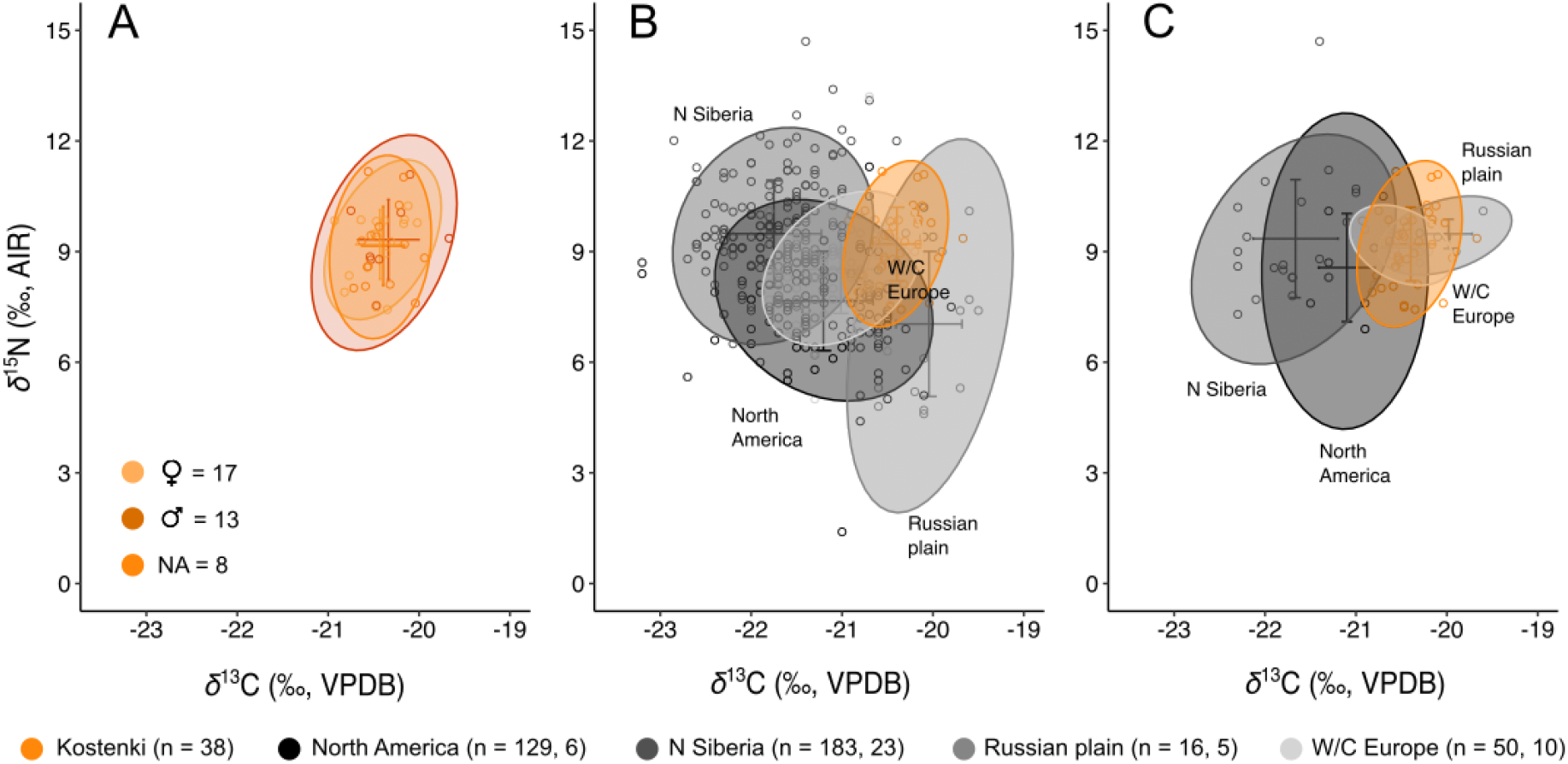
*δ*^13^C and *δ*^15^N isotopic composition of woolly mammoths. Bivariate plot of the 38 Kostenki 11-Ia woolly mammoth isotopic compositions (A) by biomolecular sex; and in the context of (B) 378 published woolly mammoth isotopic records with ages from >55,500 ^14^C yr BP (infinite) to 13,871 cal yr BP; and (C) woolly mammoth records from the Last Glacial Maximum only (24,600-17,000 ^14^C yr BP / 28,660-20,520 cal yr BP). Sample sizes for each region are presented for both datasets, with (n = B, C). For (A), N/A indicates sex could not be determined. Ellipses represent 0.95 confidence levels. Crosses represent mean value ± s.d.

For both Hap13 and 41, we found no SNPs in the control region, indicating the sequences across individuals were indeed identical. This included the three Kostenki specimens that had >20% N. For Hap13, also the older Kostenki individual (14) shared the control region sequence, as did two of the three Polish individuals; only sequence MF579947.1 from Kraków Spadzista differed with a T in its position 16,209, where the others have a C (or N).

Hap74 had two SNPs, with G to A and T to C transitions in specimen 18232 at position 16,294-16,295 relative to specimen 18251 (determined by 13 reads). In specimen 18232, only three reads cover these positions, with one having GT and two having AC, but at the first two positions of the reads. Thus, the SNPs are likely due to ancient DNA damage, and we therefore infer that the samples likely have the same haplotype in reality.

The analysis of Hap42 revealed one variable site between individuals, with a C to T transition in specimen 18243 at position 15,609, supported by 109 reads with T, often in the middle of the reads, and one read with A; in specimen 18222, the C at position 15,609 was supported by its position in the middle of 3/3 reads. The high coverage at the site in specimen 18243, and the presence of the variable site in the middle of the reads, suggested this was a real SNP and not the result of ancient DNA damage.

Five of the six identified Kostenki haplotypes fall within woolly mammoth mitochondrial Clade 1/DE (Hap13, 41, 42, 43, 89; 11/13 individuals) (Fig. 3). Clade 1/DE has previously been found in woolly mammoths from E Beringia and Eurasia (Russia, Poland, Switzerland). The remaining haplotype falls within Clade 3/B2 (Hap74; 2/13 individuals). Clade 3/B2 has been found in woolly mammoths across Europe, Russia, Ukraine, and China (14, 36).

### Specimens with radiocarbon dates and mitogenomes

Four individuals for which mitogenomes were retrieved – representing three haplotypes Hap13, 41, and 89 – also had reliable radiocarbon dates; although we also retrieved a mitogenome for specimen 18222, the very young date of this specimen was deemed unreliable and was omitted from further analysis (Table S3).

The four individuals grouped in all three identified phases (Fig. 2). Specimen 18256 (UCIAMS-266006; Hap41; genetically identified as female) was the youngest of the two oldest samples identified at the site. Interestingly, Hap41 – unique to our data – was shared with specimen 18233, which was placed in the centre of the mammoth bone structure, and was identified as a potential male based on palaeoproteomic analysis (Fig. S3, Table S3). We did not have a radiocarbon date for specimen 18233, and thus future dating efforts are needed to place the individual in a temporal context.

Hap13 was shared by specimens 18244 (UCIAMS-266004; genetically identified as male) and 18239 (UCIAMS-266003; genetically identified as female). The two individuals sharing Hap13 did not group in the same phase; 18244 was grouped in Phase 1 and 18239 in Phase 2, and thus they are unlikely to be contemporaneous.

Specimen 18250/UCIAMS-251307, genetically identified as male, had a unique haplotype and grouped in Phase 2.

### Stable isotope analysis

We generated 38 *δ*^13^C and *δ*^15^N measurements from bone (n = 1) and dentine (n = 37) collagen of the Kostenki woolly mammoths (Fig. 4); sample 18253 did not yield enough collagen for stable isotope analysis. We combined our data from the 38 individuals with publicly available records from the species. Based on the overlap on the 95% confidence ellipses and the average estimates, we did not observe any significant differences in isotopic composition between sexes within the Kostenki 11-Ia woolly mammoths (Fig. 4A).

The published dataset comprised 378 woolly mammoths from E Beringia (n = 129), N Siberia (n = 183), Russian Plain (n = 16), and W/C Europe (n = 50), with sample ages from >55,500 ^14^C yr BP to 13,871 cal yr BP (Table S8). The average *δ*^13^C and *δ*^15^N of the Kostenki mammoths falls close to values recorded for the Russian Plain and W/C Europe (Fig. 4B). When focusing only on samples from the LGM, the average isotopic composition of the Kostenki 11-Ia woolly mammoths are also closer to Russian Plain and W/C Europe woolly mammoths (Fig. 4C).

## Discussion

We performed biomolecular analysis of 39 woolly mammoth specimens sampled from the third mammoth bone structure at Kostenki 11-Ia. We radiocarbon dated nine specimens (one specimen was dated twice, totalling ten dates), and used the ages to interpret the time frame of human activity at the site. Combining genetic and proteomic methods, we sexed 30 individuals, and further contextualised the Kostenki 11-Ia specimens within the frameworks of available mitochondrial DNA and paleoecological (*δ*^13^C and *δ*^15^N) woolly mammoth data.

### Dating human activity at the site

The 11 high-precision radiocarbon dates – which included mammoth teeth and charcoal – indicated the presence of three radiocarbon date groups, and in addition indicate two phases of human activity at the site (Fig. 2). The dates new to this study indicate the newly-discovered presence of some older mammoth material at the site, in the form of two molars that are placed in the outer ring of the structure (Fig. S2). Interestingly, the youngest of these two older samples shared a haplotype with another individual at the site, which was placed in the centre of the structure (Fig. S3). The sample was not dated, and thus we cannot place it in a temporal context.

Based on a smaller set of dates for Kostenki 11-Ia, Pryor et al. (3) reported evidence for two distinct clusters of dates at the third structure, and suggested this indicated either a single occupation phase plus a group of younger contaminated dates; or that there were two discrete occupation phases at the site. The expanded dataset presented here shows a similar pattern, whereby the high-precision radiocarbon dates can be divided into two distinct groups – Phase 1 aged 25,040 - 24,670 cal BP and Phase 2 aged 24,580 - 24,230 cal BP (Fig. 2).

The apparent hiatus between the two newly-defined phases is substantially less than that reported in Pryor et al. (3), although it remains in the order of hundreds of years. It is notable that both identified phases are represented by dates on two different materials, and include mammoth specimens and charcoal recovered from various places within the third mammoth bone structure, potentially indicating human activity in both phases. We further note that the oldest and only reliable available date for the first Kostenki 11-Ia structure (GIN-2532; 24,900 −23,100 cal yr BP) correlates broadly with the Phase 2 dates of the third structure (Fig. S1), suggesting Phase 2 may have included activity at both the first and third Kostenki 11-Ia structures, a point which warrants further investigation in future.

The discreteness of the radiocarbon dates into two phases is consistent with the discovery in 2014-2015 of a stratified area of burning at the site, which indicates two phases of activity (10). Our biomolecular data do not shed further light on this; of the individuals with radiocarbon dates and mitogenomes, two shared a haplotype, and had overlapping 95.4% age ranges, which could lend support to them being part of the same family group or matriline. However, the dates were grouped in different phases. Furthermore, Hap13 was the most prevalent haplotype – found in eight of our individuals, the available Kostenki mitogenome (14), and three Kraków Spadzista mammoths (36). The radiocarbon ages of the Hap13 individuals indicate the presence of the haplotype at Kostenki across at least 11,000 years, from 35,170 to 24,402 cal yr BP, and thus it is not an informative lineage for inferring familial relationships.

Combined, our findings indicate two interpretations of the Phase 1/Phase 2 radiocarbon dates at Kostenki 11-Ia are possible, with the evidence indicating: i) a single phase of human activity, that involved the use of already-old mammoth bones and wood hundreds of years old at the time humans brought them to site; or ii) that there genuinely were two discrete phases of activity at Kostenki 11-Ia, separated by hundreds of years. Distinguishing these possibilities will require further investigation of the site’s archaeology, especially an understanding of the stratigraphic context and material remains found there, to discern whether or not different phases of activity can be identified.

### Modes of bone accumulation at Kostenki 11-Ia

Our biomolecular sexing showed a predominance of females among the mammoth specimens at Kostenki 11-Ia (♀ = 57%, ♂ = 43%; Table S3). Genetic sexing of mammoth bones collected across a large geographic area in NE Siberia identified a majority of males (n = 98; ♀ = 31%, ♂ = 69%) (15). This was explained by putative differences in behaviour of females and males, and the social structure of woolly mammoths (e.g., 37); less experienced solitary males may have been more likely to be caught and die in natural traps, which favour preservation, and thus are overrepresented in the fossil record (15). The overrepresentation of females at Kostenki suggest the individuals were derived primarily from mammoth herds, which is supported by the presence of all age classes at the site; depending on area, the percentage of juveniles varies across the structure, from 3-4% to 10-15% of the bones (12–38). If specimens were from natural traps only, we would expect more age uniformity, with mostly subadult and adult males.

Our radiocarbon evidence indicated the Kostenki 11-Ia mammoths died across several centuries, and thus did not represent a single-family group. This was further verified by our identification of six mitochondrial haplotypes among 16 individuals (Fig. 3, Table S3). Three haplotypes were present in multiple individuals. Hap13 was found in eight of our Kostenki individuals, but our analysis shows it had a wide geographic range and had been present in Kostenki for at least the previous 11,000 years. Two haplotypes unique to our samples were each present in two individuals, which could indicate the individuals were related, as the mitochondrial genome is passed from mother to offspring. However, only one of the samples (18256; UCIAMS-266006; Hap41; female) was radiocarbon dated, and we do not have age data of the corresponding sample (18233) to place it in a temporal context. For the other haplotype pair (Hap42), specimen 18222 yielded a too-young date and could not be re-dated due to lacking collagen, and hence we will not be able to assess the temporal context of both these individuals. Thus, we are unable to evidence further if the mammoths sharing the unique Kostenki haplotypes are from the same family group or matriline, which would be supported by similar radiocarbon ages.

Only a small number of the bones excavated in the structure were articulated, primarily vertebral bones (10), suggesting the mammoths did not die on site. This pattern differs from the woolly mammoths at the Sevsk family group site, which were partially or completely articulated (21). Thus, we suggest the woolly mammoths at Kostenki 11-Ia died off-site, and were harvested and moved to the structure. Indeed, the lower fraction of juveniles present at Kostenki 11-Ia relative to bone beds of a herd or a whole population – where the number of juveniles comprise ∼40% of the individuals (39) – may suggest the preferential acquisition of adult individuals.

Our analysis identified two new dates measured on mammoth specimens from the third structure that were much older than the rest; UCIAMS266007 (18257; genetically sexed as male) and UCIAMS266006 (18256; genetically sexed as female), aged 25,460 and 25,210 cal yr BP (median ages), respectively (Fig. 2). The mandibles were positioned along the outer edge of the ring of bones making up the eastern wall of the structure (Fig. S2). The ages were too old to be modelled with Phase 1 dates, being 0-1,300 years older, and 200-1,500 years older than Phase 2 dates (95.4% certainty, Fig. 2).

We are confident the two older dates do not conform with the main phases of human occupation at the site, due to the strength of evidence constraining the two inferred phases (i.e., number of individual dates) and, in addition, the extended period of human activity suggested by their inclusion is not supported by the archeology of the site (3). We are also confident that the two older mandibles were not redeposited from older sedimentary layers at Kostenki 11-Ia due to their position within the ring of bones defining the structure. Redeposition has been reported in the mammoth bone assemblage of Achchagyi–Allaikha, where a specimen ∼24,000 yr older than any other mammoth radiocarbon dates at the site was interpreted as a redeposition from an older sedimentary layer (40). However, the evident stratigraphic integrity of Kostenki 11-Ia argues against this.

Rather, we interpret our dates as evidence that the two oldest-dated mandibles were scavenged from long-dead carcasses of mammoth, and brought to Kostenki 11-1a as ‘fossil’ raw bone material by the humans that built the structure. The older material may have been scavenged from other Kostenki sites, where both natural and human-created mammoth bone beds have been discovered (e.g., in Kostenki 5-layer II, a natural bone bed with dates from ∼28,000 - 24,500 cal years BP; reviewed in 41).

Based on the available data, we are unable to elucidate if the Kostenki 11-Ia mammoths were also derived from active human hunting. Human exploitation of mammoth bones and ivory from natural accumulations has been documented elsewhere on the basis of radiocarbon dating evidence (e.g., 42), and has been widely discussed at a theoretical level in the context of the circular mammoth bone features (e.g., 43-44). However, to our knowledge, our analysis provides what may be the first direct evidence indicating the use of scavenged skeletal material in the construction of a circular mammoth bone feature, and is thus a significant finding that adds to our understanding of how these sites were created.

### Phylogeography and palaeoecology of the Kostenki 11-Ia mammoths

We analysed the 16 retrieved mitochondrial genomes, representing six haplotypes, in the context of available data to gain insights into the phylogeography of the Kostenki 11-Ia mammoths. The phylogeny of mammoth mitochondrial DNA indicates three main clades (Clade 1-Clade 3), which can be further subdivided into several subclades that are somewhat geographically distinct (e.g., 14, 23, 36, 45).

The majority of our mammoth samples fall within Clade 1/DE. The only available mitogenome from Kostenki (unknown site) retrieved from a pre-LGM specimen (14) also grouped in this clade. One haplotype unique to Kostenki and found in two of our individuals grouped in Clade 3/B2 (Fig. 3). Our findings are in agreement with the known distribution of the two clades. Clade 1/DE extended from E Beringia to western central Europe (e.g., 14, 36). This clade has been recovered in mammoths from E Beringia ranging in age from ∼37 to ∼16 thousand years ago (ka). In Siberia (including NE Siberia and southern central Siberia) there was a long-term distribution of this clade, and it has been sequenced in mammoths aged >40 ka to ∼4 ka. The youngest specimens from this clade (∼7-4 ka) were all from NE Siberia. In western and central Europe, this clade has been found in LGM samples from Poland and in a post-LGM mammoth from Switzerland (Fellows Yates et al. 2017). Clade 1/DE has been reported in mammoths across Europe, Russia, Ukraine, and China (14).

Our study provides, to our knowledge, the first attempt to investigate differences in foraging ecology between females and males in woolly mammoths, and indeed in any Late Pleistocene megafaunal remains. Foraging differences between sexes, based on behavioural observations and measurements of elephant feeding impact on the tree canopy, have been reported for African elephants, and are influenced by body size, reproductive strategies, and social structure (46). Foraging differences between females and males have also been described for Asian elephants (47). Based on bone and dentine collagen *δ*^13^C and *δ*^15^N, we did not detect differences in stable isotope composition between sexes (Fig. 4). This finding may reflect a lack of resource partitioning between sexes, albeit the coarse nature of our data is such that fine-scale differences cannot be identified. Bone and dentine collagen *δ*^13^C and *δ*^15^N provides insights into long-term foraging averaged over years to decades, but does not provide information on what species comprise the diet (e.g., 48). However, when comparing *δ*^13^C and *δ*^15^N among species, they can be used to identify differences in foraging or how environmental factors differentially affect species foraging ecology. For instance, based on *δ*^15^N, mammoths are known to present a distinct isotopic composition compared to other herbivores (e.g., horse), which may reflect different plant type or plant part preferences (49).

Our sample sizes of the two sexes differ, and median calibrated ages of the radiocarbon dates of our samples span up to 1,070 years, which may mask differences and not fully capture mammoth foraging strategies. The *δ*^13^C and *δ*^15^N data obtained for the Kostenki 11-Ia mammoths show similar average values to mammoths from the Russian plain (Eurasia lower latitudes <60°N) and W/C Europe, when focusing on the LGM period (Fig. 4). Mammoths are known to show regional differences in isotopic compositions (e.g., 17) and thus the isotopic compositions of the Kostenki mammoth agree with the geographic location of the site.

Our study provides new insight into the archeological context of the mammoth bone structures of the central European Plain, and the Palaeolithic humans associated with them. The eight new reliable radiocarbon dates presented in this study confirmed the third structure at Kostenki 11-Ia is among the oldest circular mammoth bone structures yet discovered. Furthermore, the dates indicate human activity at the site either spanned several centuries, potentially occurring in two discrete phases, or involved collection of already-ancient bones and wood from across the landscape. In combination with the archaeology of the site, our data suggest the mammoth bones were acquired off-site, including at least some degree of scavenging from bone beds and opportunistic finds of long-dead individuals. Our study also provides novel insights on sex differences in the foraging ecology of woolly mammoths.

## Materials and Methods

### Experimental design

We performed biomolecular analysis of 39 woolly mammoth specimens sampled from the third mammoth bone structure at Kostenki 11-Ia. We radiocarbon dated nine specimens (one specimen was dated twice, totalling ten dates), and used the ages to interpret the time frame of human activity at the site. Combining ancient DNA and palaeoproteomic methods, we sexed the mammoth individuals, and used the data to identify the ratio of females and males at the site. By integrating biomolecular sexing with stable *δ*^13^C and *δ*^15^N isotope analysis of the same specimens, we investigated differences in isotopically-differentiated resource use by females and males. Mitochondrial genomes and paleoecological (*δ*^13^C and *δ*^15^N) data from the Kostenki 11-Ia individuals were contextualised within the frameworks of available data from other late Quaternary woolly mammoths.

### Mammoth specimens

A minimum of 51 mandibles and 64 crania from mammoths had been excavated at the third mammoth bone structure at Kostenki 11-Ia by the end of 2015 (Voronezh, Russia; Fig. 1D) (12). E.D.L. and A.M. visited the site in August 2015 to collect mammoth faunal material for ancient biomolecular work. At that time, 40 mandibles had been unearthed and could be identified in the structure. To ensure that each sample represented a unique individual, protruding molars of the left mandible were sampled for 37 individuals. For two individuals, where the mandible was deposited upside down, direct access to the tooth was not possible and instead bone from the left mandible was sampled (Table S3). For ease of reference, we have excluded the prefix of ‘CGG_1_0’ from the sample IDs, and only use the last five digits in the text, e.g. sample CGG_1_018221 is designated 18221. As the excavation was still ongoing, we cannot match the sampled individuals with the final distribution maps of the site.

The sampled mandibles varied in size and most likely represent both adults and juveniles (Fig. 1C). However, no data are available on the relative size of the individuals sampled. In 2016, 44 (out of 45 unearthed open mandibles) were measured, revealing that some were significantly smaller. However, a more comprehensive analysis is still required to determine whether these smaller specimens are juvenile mammoths. In addition, in depth analysis on the anthropogenic modifications on the mammoth bones is ongoing.

### Radiocarbon dating

We radiocarbon dated nine specimens at the Keck Carbon Cycle AMS Facility (Earth System Science Department, UC Irvine) (Table S3). Subsampling of the specimens was conducted at the ancient DNA clean lab facilities at Globe Institute (University of Copenhagen), and collagen was extracted at Trent University (Beaumont et al. 2010). 100-200 mg of bone/dentine chunks were demineralized in 0.5 M HCl for approximately 24 h. Samples were then rinsed to neutrality with ultrapure water. Any samples exhibiting a dark discoloration were treated with 0.1 M NaOH for successive 20 min treatments until no colour change was observed in the solution. Samples were rinsed to neutrality with ultrapure water. All demineralized samples were then placed in 4 mL of 0.01 M HCl and the collagen was solubilized at 65°C for 36 h. The resulting solution containing the collagen was then filtered using 30 kDa centrifugal filters (Centriprep, Millipore) that were cleaned according to Beaumont et al. 2010. The >30 kDa portion was then transferred to a glass vial, frozen and lyophilized.

We calibrated the radiocarbon dates in OxCal v.4.4 (33) using the IntCal20 (50) calibration curve. Two of the dates (UCIAMS-251304 from specimen 18222; UCIAMS-251305 from specimen 18241) were younger than our expectations based on the archeological chronology. Therefore, we re-sampled the two specimens and carried out new collagen extractions for an additional round of dating. However, one specimen (18222) did not yield enough collagen to provide a second date, resulting in a final dataset of ten new radiocarbon dates. With each batch of samples for which collagen was prepared for radiocarbon dating, one or more aliquots of a sample with an infinite radiocarbon age (Hollis Mine mammoth, FmC = 0.0031 ± 0.0002) (51) and a secondary standard of known age (Umingmak whale, 7325 ± 40 ^14^C yr BP) (52) were prepared in the same manner as the samples, and radiocarbon dated. These standards returned radiocarbon ages very similar to their long-term measured values: FmC = 0.0031 ± 0.0009 for the Hollis Mine Mammoth and 7340 ± 20 ^14^C yr BP for the Umingmak whale.

We analysed the total of ten dates (including the two dates generated for specimen 18241) with 14 available radiocarbon dates from Kostenki 11-Ia from the first and third structures at the site (Table S4), to create an overview of the chronology of this site, using in OxCal v.4.4 (33) and IntCal20 (34). Available dates comprise four charcoal dates, and ten mammoth bone dates (3, 9, 53–55). Further analysis was then conducted on a reduced dataset, excluding erroneous dates and also dates measured with low precision using OxCal v.4.4 (33), with the IntCal20 calibration curve (34). This reduced dataset comprised eight new radiocarbon dates reported here, and three previously measured charcoal dates. Further details are provided in the Supplementary Text.

### Ancient DNA extraction and sequencing

We drilled 50-70 mg of bone or dentine powder from each mammoth specimen. DNA extractions were carried out following two different protocols: a silica-powder based method and a silica column method (Table S5). The silica-powder method is based on the protocol published in Rohland and Hofreiter (56) with some modifications following Allentoft et al. (57). We included a pre-digestion and a modified binding buffer as described in Allentoft et al. 2015. The modified binding buffer was prepared in bulk by mixing 500 mL Qiagen buffer PB with 9 ml sodium acetate (5M) and 1.25 mL sodium chloride (5M). After mixing the pH was checked and corrected until reaching a value between 4-5.

The second DNA extraction protocol used a modified version of a silica-column based protocol (58). Dentine/bone powder was incubated overnight at 37°C with constant rotation in 1 mL of extraction buffer. After the overnight incubation, the supernatant was added to a 30 KDa Amicon® Ultra-4. The sample was centrifuged at 4,000 rpm, until the supernatant was concentrated down to 70 μL. The concentrate was combined with 10x modified Qiagen PB buffer as described in (57) and purified using Monarch columns (NEB). After DNA binding to the Monarch columns, we performed two washes with Qiagen PE. DNA elution was performed in two steps; for every step, we added 25 μL of Qiagen EB buffer to the Monarch column, incubated for 5 min at room temperature, and centrifuged at 13,000 rpm (max speed) for 1 min.

Some of the DNA extracts were treated with Thermolabile USER II enzyme (NEB) prior to library build (Table S5). For each sample, the USER reaction was performed in 16 μL total volume, with 2.4 μL of the Thermolabile USER II enzyme and 13.6 μL of each extract, and incubation time of 3 h at 37°C. USER treated DNA extracts were purified using Monarch columns (NEB).

We used two different library build protocols: a double stranded DNA (dsDNA) (59) and a single stranded DNA (60) method (Table S5). For the dsDNA library build, we used the protocol described in Meyer and Kircher (59) with the following modifications: our reaction volume was 25 μL, the initial DNA fragmentation was not performed, and MinElute kit (Qiagen) was used for the purification steps. The ssDNA libraries were built following the procedure described in Kapp et al. (60). Only the USER treated extracts were used for the ssDNA protocol. All the libraries were double-indexed using KAPA HiFi HotStart Uracil+ReadyMix (Roche). The resulting indexed libraries were quantified on a Qubit^TM^ dsDNA HS (Invitrogen) and quality checked in the Agilent BioAnalyzer or Fragment Analyzer. Indexed libraries were shotgun sequenced on an Illumina HiSeq 4000 80 base pairs (bp) or NovaSeq 6000 150 PE.

### Bioinformatic data processing

Adapter and quality trimming with AdapterRemoval v2.2.0 (61), mapping (read alignment, PCR duplicate removal, and indel realignment), and mapDamage v2.0.6 analyses (62) of the shotgun data were performed using the PALEOMIX pipeline v1.3.6 (63). For the samples sequenced in Illumina NovaSeq 150 PE, we only used the R1 in our analysis. Due to the short length of the recovered DNA fragments, we would not expect to recover extra data by analysing the R2. Mapping was performed using the BWA-aln v0.7.17 (64) with seed length disabled to improve mapping efficiency (65). Reads shorter than 30 bp were discarded during adaptor trimming and reads showing mapping qualities less than 30 were also excluded. For mapping, we used the nuclear genome assembly of the African savannah elephant (*Loxodonta africana*; LoxAfr4) and a woolly mammoth mitochondrial genome (mitogenome) reference (GenBank: DQ188829.2).

We generated mitogenome consensus sequences for the three samples that had >1,000 reads mapping to the mitogenome reference (Tables S3, S5). We generated the consensus sequences using Geneious R11 (66) using the strict criteria requiring more than 50% of reads for each base to match for bases with a coverage >3x. Bases with coverage <3x or ambiguous bases were called as undetermined (Ns).

### Mitochondrial genome capture

To increase the number of mitochondrial sequences, we performed mitochondrial enrichment by target capture on the 20 ssDNA libraries using a myBaits (MYcroarray/Arbor Biosciences) custom-design mitochondrial genome array, which included baits designed based on a mammoth mitogenome reference (DQ188829.2).

The myBaits v5.01 High Sensitivity protocol was used with hybridization at 55°C at all relevant steps, with the only deviation from the protocol being that only one round of enrichment was done, with a ratio of 1:4 baits:water in the hybridization mix. Post-capture amplification was performed using NEBNext Q5U Master Mix (New England Biolabs) as follows: 98°C for 2 min, 18 cycles of 98°C for 20 s, 60°C for 30 s, and 72°C for 45 s, with a final extension step of 72°C for 5 min. The captured libraries were pooled equimolar and sequenced on an Illumina NovaSeqX PE150 bp. The data processing of these reads was performed with PALEOMIX as above, except that reads were only mapped to the woolly mammoth reference mitogenome and only collapsed reads were used, as non-collapsed paired-end reads are more likely to be contaminants.

Consensus sequences were generated in Geneious as above for the 14 samples with >900 reads mapping to the reference (Table S5).

### Genetic sex determination

We performed genetic sex determination of the 23 mammoth specimens that had >5,000 reads (67) from the shotgun sequencing mapping to the nuclear genome assembly of the African savannah elephant. Sex determination was performed as described in (15). We estimated the ratio of the number of reads mapping to sex chromosome X *versus* an autosome. We used chromosome X (ChrX) and chromosome 8 (Chr8), which are of comparable size. As female mammals have two copies of chromosome X, and males carry only one copy, we expected ChrX:Chr8 ratios of ∼0.5 for males and ∼1 for females. The number of reads mapping to ChrX and Chr8 was obtained using samtools idxstats (64). To correct for chromosome size differences, the number of mapped reads was normalised by the length of the chromosomes (Table S5). Samples were determined as males if ChrX:Chr8 <=0.7, females ChrX:Chr8 >=0.8, and undetermined (N.A.) if ChrX:Chr8 0.7-0.8.

### Proteomic sex determination

Dental powders of 21 mammoths were provided for palaeoproteomic dental enamel sex assignment (Table S7). Of these specimens, three were male and three were female based on the previous ancient DNA sex assignment (see above), while the remaining 15 specimens could not be sexed based on ancient DNA analysis.

Dental enamel proteomes are dominated by peptides resulting from the *in vivo* hydrolysis of amelogenin. Amelogenin isoforms located on the X-chromosome and Y-chromosome have different amino acid sequences for some mammalian clades, allowing the proteomic sex assignment of, for example, human individuals through the observation of peptides uniquely matching to amelogenin-Y (AMELY) as male individuals. In contrast, the absence of AMELY-specific peptides combined with an abundance of AMELX-specific peptides is often taken to suggest a female sex assignment (68–70). Since dental enamel proteomes preserve over longer periods of time than ancient DNA, the proteomic determination of genetic sex provides an alternative molecular approach to study sex-based differences in hunter-gather ecology and Pleistocene fauna ecologies.

These enamel powders were immersed in 1 mL of 5% hydrochloric acid (HCl) and placed at 4°C overnight. The following day, samples were centrifuged for 10 min at 3000 *g* and supernatants collected into new tubes (labelled S1). Another mL of HCl was added for another 12 hours. After the last 12 hours, the samples were not completely demineralised and were placed on a carousel at room temperature to ensure complete demineralisation. After the additional 12 hours, the samples were demineralised, and were centrifuged using the same parameters as before. The supernatants were collected into new tubes (labelled S2) before being placed at −18°C until further steps. Each extract was then loaded on an individual Evotip (Evosep, Odense, Denmark) as follows: 500 μL of S1 and 500 μL of S2 were first combined in S3 tubes; S3 solutions were acidified with the addition of 1 μL of TFA (100%) and finally, 2×200 μL of S3 solutions were loaded on the corresponding Evotip following the standard recommended protocol (71). An extraction blank was performed alongside the samples, and analysed using liquid chromatography coupled to tandem mass spectrometry (LC-MS/MS). This blank remained empty of any protein matches to dental enamel proteins, including an absence of albumin and collagen type I (COL1) peptides.

Extracts were first separated by liquid chromatography on an Evosep One (Evosep, Odense, Denmark) using the 60SPD method with a gradient of 21 min for a total cycle of 24 min. Separation was conducted using a polymicro flexible fused silica capillary tubing (150 µm inner diameter, 16 cm long) home-pulled and was packed with C18 bonded silica particles (1.9 µm diameter, ReproSil-Pur, C18-AQ, Dr. Maisch, Germany) with mobile phases consisting of A: 5% acetonitrile and 0.1% formic acid in H_2_O and B: 0.1% formic acid in H_2_O at a flow-rate of 2 µL/min. The mass spectrometer, an Exploris 480 (Thermo Fisher Scientific) was set on data dependent acquisition mode with a first MS1 scan at resolution of 60,000 on the *m/z* range 350-1,400. The twelve most intense monoisotopic precursors were selected, and were then dynamically excluded after one appearance with their isotopes (± 20 ppm) for 20 seconds. The selected peptides were acquired on MS2 with the Orbitrap with a resolving power of 15,000, HCD set at 30%, quadrupole isolation width of 1.3 *m/z* and a first *m/z* of 120.

In the absence of available amelogenin sequences for mammoths, we built a protein reference database containing entries for 12 proteins (COL1A1, COL1A2, COL17A1, AMELX, AMELY, TUFT1, AMBN, AMLT, ALB, ENAM, ODAM, and MMP20) for *Elephas maximus* and *Loxodonta africana* derived from Genbank and UniProt. The database included 16 amelogenin isoforms across both taxa, including a single, nearly complete AMELY sequence. Proteomic data analysis was conducted in Maxquant v2.1.3.0 using unspecific digestion settings, allowing for a maximum of 5 variable PTMs (deamidation (NQ), phosphorylation (STY), and oxidation (MP)). Peptides were allowed to be between 7 and 30 amino acids, with a minimum score of 40, and filtered up to 1% FDR at peptide and protein level. Other settings were left as default. Based on the results of the genetically sexed female and male mammoth individuals, minimum peptide thresholds were determined to allow the confident identification of female individuals (based on reaching a minimum number of 15AMELX-specific peptides) and male individuals (based on reaching a minimum number of 2 unique AMELY-specific peptides). These thresholds may differ depending on mass spectrometry data acquisition settings, extraction method, or data analysis approach, and are therefore not general recommendations for future studies.

### Mitochondrial phylogeography

To determine which known genetic group/s (clade/s) the Kostenki mitogenomes belong to, we compiled 166 mammoth mitogenomes available in GenBank (data downloaded on 07-05-2021). Those shorter than 16,450 bp or that had >20% N bases were excluded from downstream analyses, leaving a reference panel of 147 mitogenomes (Table S6). Of the 16 Kostenki mitogenomes generated, three had >20% N bases (18225, 18236, and 18255) and were initially excluded from the phylogeographic analysis. Thus, a total of 160 mitogenomes were aligned using the MAFFT (72) plugin v1.5.0 in Geneious. Due to potential misalignments and missing data, we removed the d-loop from the final alignment.

To estimate the number of haplotypes (unique sequences) present in the data we used Mesquite v3.81 (73) to convert all uncertainties (e.g. ambiguous bases) in the alignment to missing data, and used this alignment in DnaSP v6.12 (74) to assign sequences to haplotypes by generating a haplotype data file. Sites with gaps or missing data were not considered.

To investigate the evolutionary relationships of the haplotypes, we used the same alignment from Mesquite in PopART v1.7 (75) to generate a median-joining haplotype network. For the three Kostenki mitogenomes that had >20% N bases, we performed the haplotype assignment individually because the increased amount of missing data that the N bases introduced into the alignment would otherwise have reduced the number of haplotypes and the overall power of the analysis. We nonetheless wanted to determine their relationship to the reference panel and the other Kostenki mitogenomes. Thus, each of the three (18225, 18236, and 18255) was aligned in turn with the other 160 mitogenomes and the haplotype analysis performed as before. Furthermore, for those Kostenki samples that shared a haplotype, we compared their control region (d-loop) sequences to further refine the haplotype sharing results. Since the control region is more variable and has a higher mutation rate, it would confirm that the mitogenomes were indeed identical if samples shared the same d-loop haplotype as well. The control region was extracted and aligned via MAFFT in Geneious and the sequences were compared as in the haplotype analysis above.

We also estimated a maximum likelihood (ML) phylogeny of the 90 haplotypes in IQ-TREE v2.2.0 (76), using the Asian elephant (*Elephas maximu*s, GenBank: EF588275.2) as an outgroup. The mitogenome with the least missing data for each haplotype was selected and the sequences were again aligned with MAFFT in Geneious, the d-loop was removed, and uncertainties changed to missing data in Mesquite. IQ-TREE was run with automatic model testing and selection (-m MFP) (77), 1,000 ultrafast bootstrap replicates (-B 1000 -alrt 1000) (78), and two independent runs (--runs 2). The consensus tree was visualised using Figtree v1.4.4 (79) and further edited for visual clarity in Inkscape v1.3.2 (www.inkscape.org).

### Stable isotope analysis

A tooth or bone fragment of 100-300 mg was cut from each sample using an Ultimate XL-D micromotor. For mammal juveniles, tooth and bone can yield higher *δ*^15^N values compared to adults due to suckling as lactate is ^15^N-enriched, which results in an enrichment in *δ*^15^N in the developing tissues (80). Some of the mandibles sampled at Kostenki 11-Ia were of smaller size, thus we assume they belonged to juveniles. In addition, for the tooth, we did not target a specific layer, but rather cut a random sample consisting of several layers, therefore reducing the influence of the suckling effect for those mandibles putatively belonging to juveniles. Each fragment was then crushed into smaller pieces using a Plattner mortar and a pestle. The crushed fragments were immersed in 9 mL of 0.5 M hydrochloric (HCl) acid at room temperature for demineralization. Throughout the demineralization treatment, the samples were agitated on an orbital shaker. Samples remained in acid between 17-30 h and were removed from the demineralization solution when the fragment was soft and/or floating in solution. Immediately upon removal from the demineralization solution, each sample was rinsed four times in 10 mL of Type I water (resistivity >18.2 MΩ cm).

Following demineralization, the samples were solubilized in 3.5 mL of 0.01 M HCl at 75°C for 36 h. The samples were centrifuged to precipitate the insoluble material, and the collagen suspended in solution was transferred into a glass vial and frozen for 24 h. Once frozen, the collagen samples were lyophilized for 48 h. Dried collagen weighing between 0.5-0.6 mg was transferred into tin capsules for stable isotope and elemental analysis. These analyses were performed at the Trent University Water Quality Centre using a Nu Horizon continuous flow isotope ratio mass spectrometer paired with a EuroVector EA 300 elemental analyzer. The stable isotope results were calibrated using Vienna Pee Dee Belemnite (VPDB) for *δ*^13^C and AIR (ambient inhalable reservoir) for *δ*^15^N. International reference standards USGS40 and USGS66 were used to perform these calibrations. In-house laboratory standards SRM-1 (caribou bone collagen), SRM-2 (walrus bone collagen), and SRM-14 (polar bear bone collagen) were used to monitor analytical accuracy and precision of the analyses.

To investigate potential differences between the sexes in their isotopic niche, we combined *δ*^13^C and *δ*^15^N for the two sexes in RStudio using ggplot (81) to estimate the 95% confidence ellipses for each dataset, and the average and standard deviation. Due to the limited number of records for each of the sexes, statistical tests were not further performed.

To contextualise the data within a spatiotemporal framework, we compiled available *δ*^13^C and *δ*^15^N records for woolly mammoths from bone and dentine collagen. We used Web of Science to perform a literature search with the terms “woolly mammoth” and “stable isotope”, and “*Mammuthus primigenius*” and “stable isotope” (data search 25/08/2021). We recovered isotope records for 378 woolly mammoths (Table S8). The data were divided into four geographical regions: Eastern Beringia (E Beringia; Yukon and Alaska; n = 129), Northern Siberia (N Siberia; Chukotka, Yakutia, and Taymyr; n = 183), Russian Plain (n = 16), and Western and Central Europe (W/C Europe; n = 50). The 95% confidence ellipses, average, and standard deviation for each region were estimated as indicated above.

Kostenki 11-Ia has been dated to the Last Glacial Maximum (LGM; 24,600-17,000 ^14^C yr BP / 28,660-20,520 cal yr BP) (82). Therefore, to reduce biases due to temporal variation in isotopic composition caused by climatic and vegetation changes, we also performed a comparison limited to woolly mammoth LGM records (n = 44 published records). Number of records used for this analysis was: E Beringia n = 6, N Siberia n = 23, Russian Plain n = 5, and W/C Europe n = 10, in addition to the new records generated from Kostenki.

## Acknowledgements

We thank Eske Willerslev for his help in facilitating the field sampling campaign to Kostenki-11 by EDL and AM in August 2015, and Michelle Christel Larsen for her contribution to DNA data generation.

## Funding

This work was funded by the Villum Foundation Young Investigator Programme grant no. 13151 to EDL and an NSERC Discovery Grant (2020–04740) to PS. FW is supported through funding from the European Research Council (ERC) under the European Union’s Horizon 2020 research and innovation programme, grant agreement no. 948365. LLM is supported by a Marie Skłodowska-Curie postdoctoral fellowship funded by the European Union’s Horizon 2020 research and innovation programme (grant agreement No 101062449).

## Author contributions

Conceptualization: ARI, EDL

Methodology: ARI, AJEP, EDL

Investigation: ARI, DdJ, AM, RK, AD, LLM, GT, TW, MAT, PS

Formal analysis: ARI, AJEP, FW, PS

Visualisation: ARI, AJEP, DdJ, EDL

Supervision: EDL

Resources: EDL

Funding acquisition: EDL

Writing—original draft: ARI, AJEP, FW, PS, EDL Writing—review & editing: All co-authors

## Data and materials availability

Bam files with the unique reads mapping to ChrX and Chr8 used for sex determination are available in the SRA under the BioProject PRJNA897816. Mitogenomes are available in GenBank with accession numbers OP838911-OP838913. Proteomic raw data and MaxQuant search results have been deposited to the ProteomeXchange Consortium via the PRIDE partner repository and are available using identifier PXD048461.

Reviewer login to access proteomic data (login details removed for final submission):

Username

Password: pY6jObWW

## Notes

### Competing Interest Statement

The authors have declared no competing interest.

