## Supplementary materials for "Ancient biomolecular analysis of 39 mammoth individuals from Kostenki 11-Ia elucidates Upper Palaeolithic human resource use"

Alba Rey-Iglesia *et al.*

**This PDF file includes:**

Supplementary Text
Figs. S1 to S4
Tables S1 to S8 legends (tables uploaded as spreadsheets)
References (83 to 125)

**Supplementary Text**

The Kostenki 11-Ia radiocarbon dates

We set out to model the phases of activity at Kostenki 11-Ia by analysing our new and all previously published radiocarbon dates for this site. Of our 10 new radiocarbon dates (Table S3), eight were deemed reliable, as is further described in the main text. We compared these eight dates against all published dates (n=14) from the first and third mammoth bone structures (3,9,53-55; Fig. 3A); these were also recalibrated in OxCal as described in the main text (Table S4).

Our comparison identified six previously published dates that had ages (sometimes several) thousands of years younger than the oldest dates for the layer, which based on the archeological evidence at the site are obviously too young (3). Problems with dates of Palaeolithic age measured in the 1980s and 1990s without ultrafiltration or modern pre-treatment methods, like most of the six younger dates, are very common, and incomplete removal of contamination still remains a serious problem today, including at Kostenki sites (e.g. 30-32, 83). We therefore excluded these six erroneous dates from further consideration, leaving a preliminary dataset of eight new dates and eight previously measured dates for analysis (Table S4). These 16 dates have median ages spanning 21,530 to 23,520 cal years BP, constricting human activity at Kostenki 11-Ia to a window of approximately 2,000 years.

While such knowledge is useful, it is important to note that contextual and stratigraphic evidence indicates that human activity at the third mammoth bone structure actually occurred over a much briefer period or several periods, lasting a few years or decades at most (3). Evidence for this includes minimal carnivore and weathering damage to the majority of bones, despite the fact that some elements (primarily vertebral bones) (10) were articulated and therefore at least partly fleshed when they were deposited; and a relative paucity of domestic occupation debris, absence of a clear 'living floor', or any other archaeological evidence indicating that the site was used by humans for whatever purposes for an extended period of time (3).

We further note that the radiocarbon dataset for Kostenki 11-Ia includes measurements made in four different radiocarbon laboratories (GIN, NSKA, CURL, UCIAMS), to substantially different levels of precision (Fig. S1A, Table S4). In particular, all four retained NSKA dates were measured with substantially lower precision than the CURL and UCIAMS dates, resulting in measurement uncertainties of ± 250-520 years for the NSKA dates, compared to ± 80-160 years for the latter two labs. This results in much wider probability distribution ranges for the calibrated NSKA dates and, consequently, low-precision estimates of the age of human activity at the site. The same is true for GIN-2532, measured on the first mammoth bone structure at Kostenki 11-Ia.

When NSKA-885 was first reported – in the context of a much smaller radiocarbon dating dataset – it was retained for analysis and used to discuss possible phasing at the site (3). Here, in the context of a much larger dataset, NSKA-885 overlaps only with the youngest probability range of dates measured with low precision and very large uncertainties (i.e. NSKA-889 and GIN-2532), while showing virtually no overlap with any of the more precise CURL and

UCIAMS dates (Fig. S1B). This pattern very likely indicates incomplete removal of contamination in NSKA-885, meaning it is too young.

As already discussed, contamination has proved to be a recurrent problem with dating this particular site, despite the application of various pre-treatment methods in different labs, and also demonstrably affected two of the new initial dates reported in this study (Table S3). It is only with the addition of the eight more dates reported here that it has become possible to also recognise the likely contamination problem with NSKA-885. To refine our dating model for the site, we therefore excluded all retained NSKA dates (n = 4), which all have large measurement uncertainties, and concentrated only on dates from the third mammoth bone structure, which is the focus of this study (thus also removing GIN-2532 from the first structure). This left 11 dates, all from CURL or UCIAMS laboratories, which have unmodelled median dates spanning 25,460 - 24,390 cal years BP (Fig. 2, main text), approximately half the duration indicated by the full dataset.

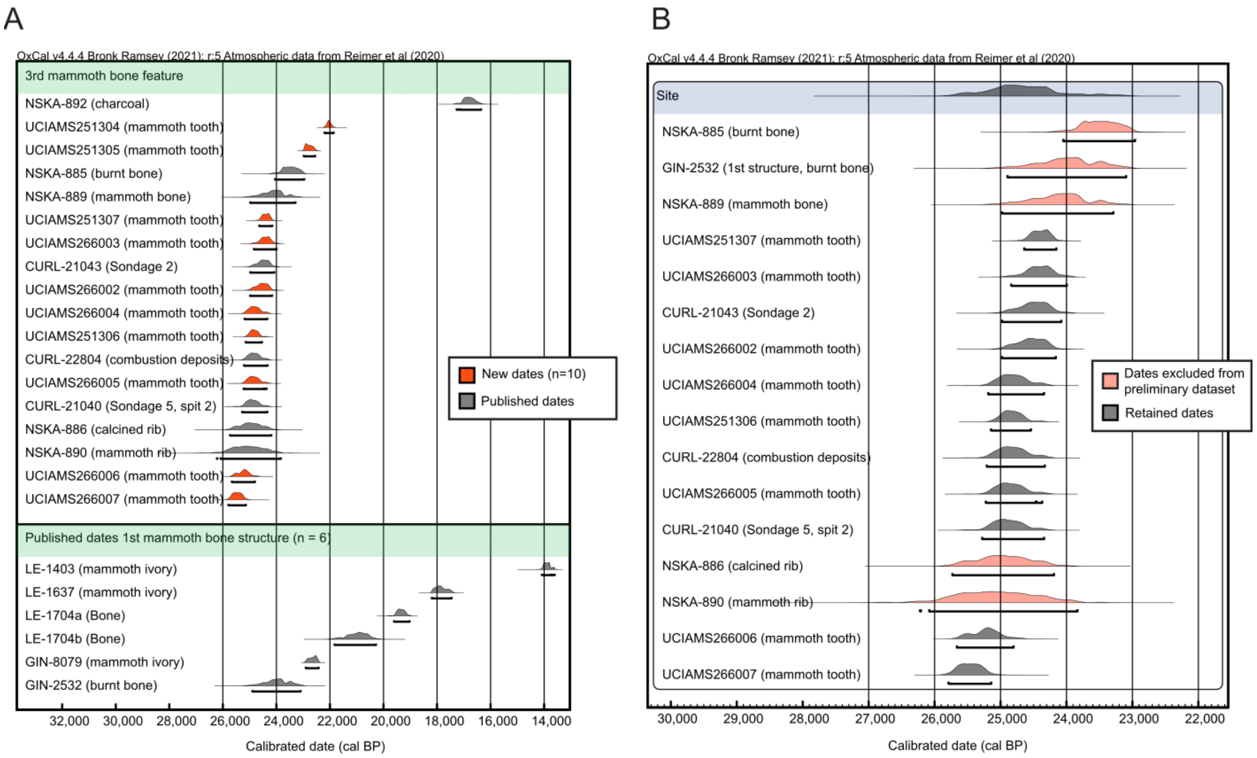


**Fig. S1.**

**Overview of the available radiocarbon dates from Kostenki 11-Ia.** (A) Calendar median year calibration of the 10 radiocarbon dates new to this study (orange) and available dates from the third (top) and first (bottom) mammoth bone structure at Kostenki 11-Ia. (B) The preliminary dataset used for interpretation, indicating dates that were eventually rejected (coral) and those retained (grey). In both panels, the substrate used for dating is indicated; full information on new and available dates is presented in Tables S3 and S4, respectively.

A B

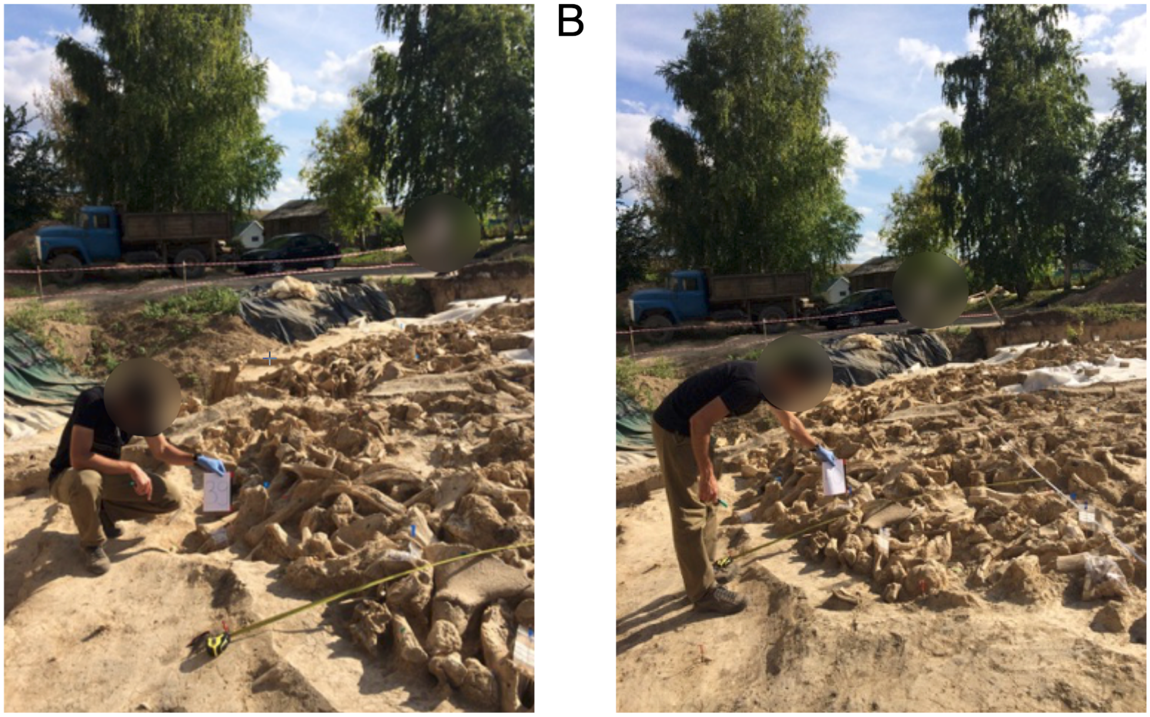

**Fig. S2.**

**The localities of the two oldest mammoth specimens obtained from the third structure at Kostenki 11-Ia.** (A) Specimen 18256 (excavation ID 39; UCIAMS-266006) and (B) Specimen 18257 (excavation ID 40; UCIAMS-266007). Both specimens were placed in the outer ring of the structure. Photo: E.D.L. during fieldwork in August 2015.

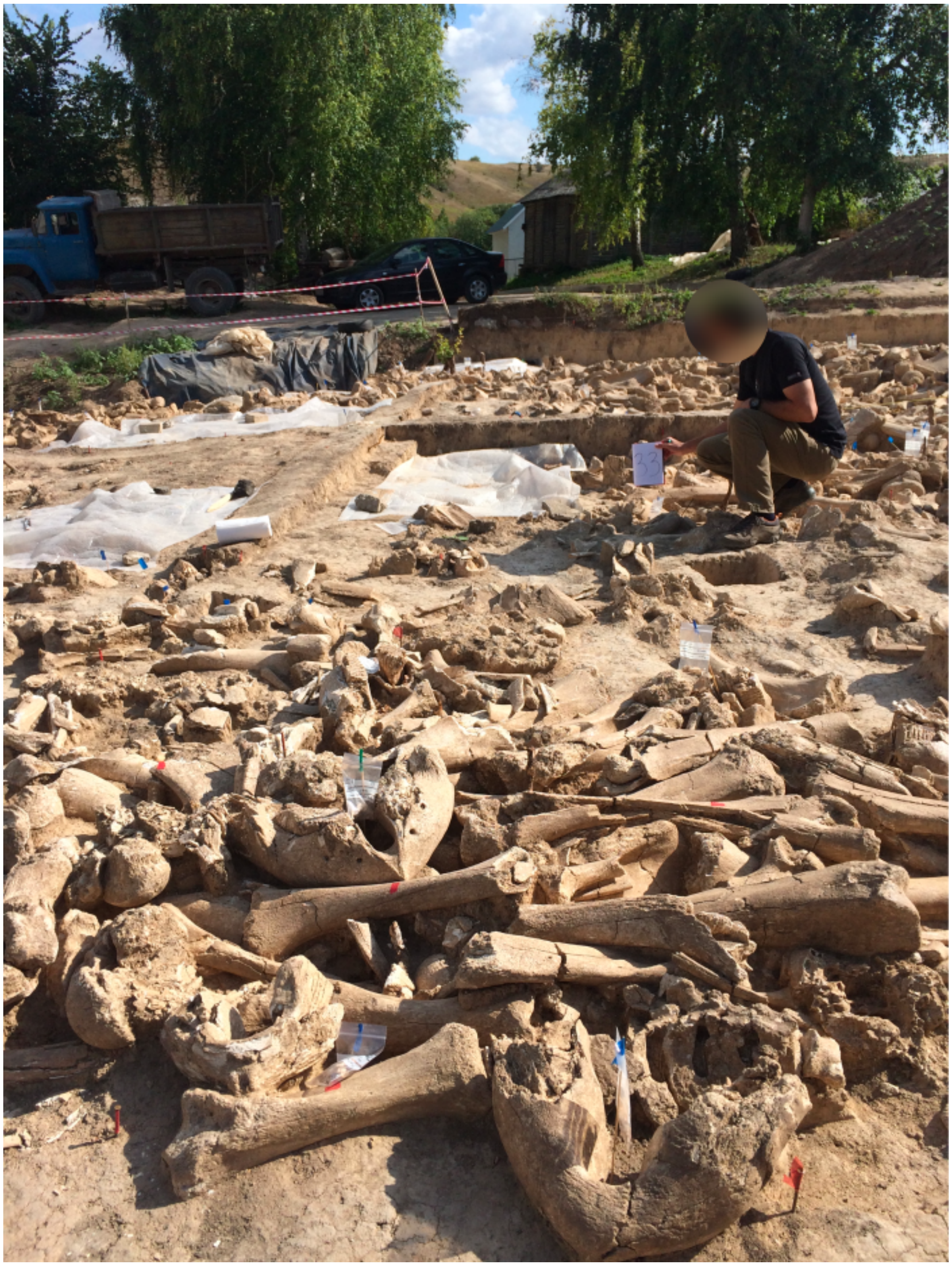


**Fig. S3.**
**The locality of specimen 18233**. The specimen (excavation ID 33) was placed in the centre of the third structure, and shared a haplotype (Hap41) – unique to Kostenki – with specimen 18256, shown in Fig S1A.

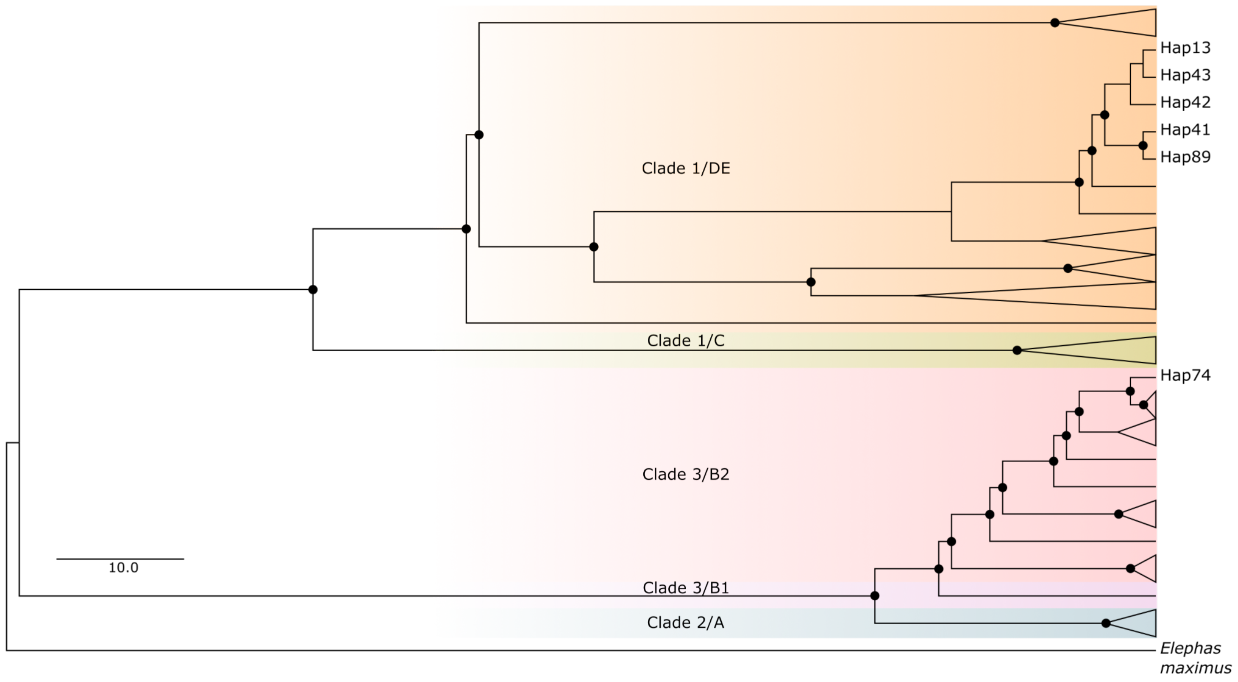


**Fig. S4.**
**Phylogenetic tree of woolly mammoth haplotypes.** Maximum likelihood phylogeny generated in IQ-TREE, with branch lengths transformed to proportional in Figtree. The alignment of 90 woolly mammoth mitogenomes and the Asian elephant (*Elephas maximus*) as outgroup contained 15,426 sites, of which 504 were parsimony informative. The six haplotypes identified in the Kostenki mammoths and discussed in the main text are indicated. Triangles represent collapsed branches. Black circles represent bootstrap support values >70%.

**Table S1.**
**Total number of woolly mammoth skeletal elements excavated at the third mammoth-bone structure at Kostenki 11-Ia.** Based on (11-12)

| Skeletal element | Left | Right | Total |
|---|---|---|---|
| Skulls | | | 64 |
| Maxilla | | | 4 |
| Zygomatic arch | | | 1 |
| Occipital condyles | | | 1 |
| Skull fragment | | | 49 |
| Hyoid bone | | | 2 |
| Lower jaw | | | 45 |
| Half of the lower jaw | 1 | 2 | 3 |
| Ascending ramus of the mandible | | | 1 |
| Symphyseal part of the mandible | | | 1 |
| Articular head of mandibular bone | | | 2 |
| Mandibular bone fragment | | | 9 |
| Atlas | | | 20 |
| Epistrophy | | | 6 |
| Cervical vertebra | | | 23 |
| Thoracic vertebra | | | 97 |
| Lumbar vertebra | | | 20 |
| Sacrum | | | 4 |
| Sacral vertebra | | | 10 |
| Tail vertebra | | | 11 |
| Atlas fragment | | | 4 |
| Odontoid process of epistrophy | | | 1 |
| Fragment of a cervical vertebra | | | 3 |
| Fragment of a thoracic vertebra | | | 7 |
| Fragment of a lumbar vertebra | | | 1 |
| Vertebral body | | | 108 |
| Vertebral body fragment | | | 15 |

| | | | |
|---|---|---|---|
| Fragments of a vertebra | | | 177 |
| Fragment of the spinous process | | | 38 |
| Vertebra | | | 22 |
| Sternum | | | 2 |
| Rib | | | 392 |
| Rib fragment | | | 309 |
| Tusk | | | 60 |
| Tusk fragment | | | 60 |
| Upper tooth | | | 4 |
| Tooth | | | 3 |
| Tooth fragment | | | 33 |
| Tooth plate | | | 66 |
| Scapula | 24 | 32 | 117 |
| Articular cavity of the scapula | | 1 | 3 |
| Blade fragment | 3 | 8 | 38 |
| Shoulder | 29 | 27 | 66 |
| Unfused articular head of the humerus | 5 | 3 | 56 |
| Unattached block of the humerus | 3 | 8 | 23 |
| Fragment of the humerus | | | 2 |
| Radial | 10 | 6 | 28 |
| Proximal end of the radius | 7 | 3 | 14 |
| Distal end of radius | 0 | 0 | 6 |
| Distal epiphysis of radius | 1 | 1 | 5 |
| Fragment of radius | | | 1 |
| Ulna | 28 | 20 | 53 |
| Unfused distal epiphysis of the ulna | 4 | 7 | 17 |
| Proximal end of ulna | 1 | 0 | 2 |
| Fragment of the distal end of the ulna | 0 | 0 | 1 |
| Hand bones | | | 50 |
| Pelvic bone | 2 | 1 | 63 |
| Articular cavity of the pelvic bone | | | 3 |

| | | | |
|---|---|---|---|
| Fragment of the pelvic bone | | 1 | 69 |
| Femoral | 32 | 18 | 63 |
| Proximal end of the femur | 2 | | 3 |
| Distal end of femur | 10 | 12 | 31 |
| Unfused articular head | 1 | | 29 |
| Unfused distal epiphysis | 4 | 4 | 8 |
| Trochanter apophysis | | | 1 |
| Fragment of the femur | | | 7 |
| Knee cap | 2 | | 12 |
| Tibia | 22 | 24 | 62 |
| Proximal end of the tibia | 1 | 3 | 8 |
| Distal end of tibia | 0 | 2 | 4 |
| Unfused proximal epiphysis of the tibia | 1 | 1 | 8 |
| Unfused distal epiphysis of the tibia | 0 | 0 | 4 |
| Fragment of the tibia | | | 4 |
| Small tibia | 1 | | 10 |
| Distal end of the fibula | | | 6 |
| Unfused distal epiphysis of the tibia | | | 4 |
| Foot bone | | | 25 |
| Navicular bone of the foot | | | 5 |
| Calcaneus | | | 23 |
| Talus | 1 | 1 | 17 |
| Medial cuneiform bone | | | 2 |
| Lateral cuneiform bone of the foot | | | 4 |
| Cuboid | | | 6 |
| Metapodial bone | | | 72 |
| Unattached distal epiphysis of the metapodial bone | | | 10 |
| Proximal end of metapodial bone | | | 1 |
| Fragment of metapadial bone | | | 2 |
| Distal end of metapodial bone | | | 2 |
| Distal epiphysis of the metapodial bone | | | 1 |

| | | | |
|---|---|---|---|
| 1 phalanx | | | 6 |
| 2 phalanx | | | 15 |
| 3 phalanx | | | 12 |
| Unfused proximal epiphysis of the phalanx | | | 1 |
| Unfused distal epiphysis of the phalanx | | | 1 |
| Phalanx fragment | | | 3 |
| Tubular bone fragment | | | 215 |
| **Total** | | | **2982** |

**Table S2.**
**Composition of fauna excavated at the third woolly mammoth-bone structure at Kostenki 11-Ia.** Based on the final excavation report from 2016 (12; original in Russian)

| **Species** | **Number of bones** | **Minimum number of individuals** |
|---|---|---|
| Woolly mammoth (*Mammuthus primigenius*) | 2982 | 64 |
| Don hare (*Lepus tanaiticus*) | 16 | 1 |
| Arctic fox (*Vulpes lagopus*) | 14 | 1 |
| Brown bear (*Ursus arctos*) | 1 | 1 |
| Cave lion (*Panthera spelaea*) | 1 | 1 |
| Horse (*Equus ferus*) | 6 | 1 |

**Table S3.**
**Sample information of the Kostenki 11-Ia woolly mammoths analysed in this study.** Information includes genetic sex identification, $\delta^{13}C$ and $\delta^{15}N$ values, and radiocarbon age; note that sample 018241 was radiocarbon dated twice. Radiocarbon dates were calibrated using IntCal20 (50). The results of the palaeoproteomic sex identification is shown in Table S7.

Attached as a spreadsheet.

**Table S4.**
**Ages of the fourteen radiocarbon dates available from the first and third mammoth bone structure at Kostenki 11-Ia.** Not all ages are considered reliable, and hence only a subset was used to estimate the phases of activity at the site. Ages were calibrated using IntCal20 (50).

Attached as a spreadsheet.

**Table S5.**
**Mapping statistics of the Kostenki mammoth DNA data to the nuclear and mitochondrial reference genomes.** Data on genetic sex determination of the individuals is included.

Attached as a spreadsheet.

**Table S6.**
**Overview of the 160 woolly mammoth mitogenome sequences analysed.** Of these, 147 were downloaded from GenBank.

Attached as a spreadsheet.

**Table S7.**
**Palaeoproteomic sex assignment.** Counts indicate the total number of unique+razor peptides assigned to collagen type I, summed for COL1A1 and COL1A2, to amelogenin, summed for AMELX and AMELY, and for AMELX and AMELY separately. The associated extraction blank did not contain any amelogenin or collagen type I peptides.

| Specimen (CGG number) | Genetic sex | COL1 unique+razor peptide count | AMEL* unique+razor peptide count | AMELX unique+razor peptide count | AMELY unique+razor peptide count | Proteomic sex assignment |
|---|---|---|---|---|---|---|
| 018221 | Female | 20 | 146 | 146 | 0 | Female |
| 018222 | Female | 75 | 20 | 20 | 0 | Female |
| 018226 | Female | 54 | 15 | 15 | 0 | Female |
| 018223 | Male | 200 | No amelogenin identified | | | |
| 018238 | Male | 0 | 107 | 96 | 11 | Male |
| 018248 | Male | 2 | 45 | 43 | 2 | Male |
| 018224 | Unknown | 2 | 105 | 97 | 8 | Male |
| 018225 | Unknown | 191 | 3 | 3 | 0 | (Female?) |
| 018229 | Unknown | 0 | 63 | 59 | 4 | Male |
| 018230 | Unknown | 18 | No amelogenin identified | | | |
| 018232 | Unknown | 1 | 137 | 126 | 11 | Male |
| 018233 | Unknown | 170 | 33 | 32 | 1 | (Male?) |
| 018234 | Unknown | 155 | No amelogenin identified | | | |
| 018235 | Unknown | 55 | 74 | 74 | 0 | Female |
| 018236 | Unknown | 76 | 5 | 5 | 0 | (Female?) |
| 018243 | Unknown | 42 | 31 | 31 | 0 | Female |
| 018244 | Unknown | 0 | 93 | 91 | 2 | Male |
| 018246 | Unknown | 194 | No amelogenin identified | | | |
| 018249 | Unknown | 28 | No amelogenin identified | | | |
| 018255 | Unknown | 0 | 122 | 121 | 1 | (Male?) |
| 018259 | Unknown | 53 | 128 | 118 | 10 | Male |

**Table S8.**

**Overview of the 378 available woolly mammoth isotopic records.** The regions used in this study are North America: Northern East Beringia, Central East Beringia; N Siberia: Taymyr (Siberia, West of 127°E), Yakutia and Chukchi (Siberia, East of 127°E); Russian Plain: Eurasia lower latitudes <60°N; W/C Europe: W/C European countries. The time bins used are pre-LGM (>24,600 $^{14}C$ years BP / 28,660 cal years BP); LGM (24,600 - 17,000 $^{14}C$ years BP / 28,660 - 20,520 cal years BP); post-LGM (<17,000 $^{14}C$ yearsBP / <20,520 cal years BP)

Attached as a spreadsheet.
